# Human-like dissociations between confidence and accuracy in convolutional neural networks

**DOI:** 10.1101/2024.02.01.578187

**Authors:** Medha Shekhar, Dobromir Rahnev

**Affiliations:** School of Psychology, Georgia Institute of Technology, Atlanta, GA

**Author notes:** **Competing Interests** The authors declared no competing interests. **Author Contributions** *MS*: Formal analysis, Investigation, Software, Data curation, Writing-Original draft preparation, Visualization; *DR*: Supervision, Validation, Funding acquisition; All authors: Methodology, Conceptualization, Writing-Reviewing and Editing. **Corresponding Author:** Medha Shekhar, School of Psychology, Georgia Institute of Technology, 654 Cherry Str NW, Atlanta, GA 30332.

**Keywords:** Artificial neural networks, confidence, perceptual decision making, visual metacognition

## Abstract

Prior research has shown that manipulating stimulus energy by changing both stimulus contrast and variability results in confidence-accuracy dissociations in humans. Specifically, even when performance is matched, higher stimulus energy leads to higher confidence. The most common explanation for this effect is the positive evidence heuristic where confidence neglects evidence that disconfirms the choice. However, an alternative explanation is the signal-and-variance-increase hypothesis, according to which these dissociations arise from low-level changes in the separation and variance of perceptual representations. Because artificial neural networks lack built-in confidence heuristics, they can serve as a test for the necessity of confidence heuristics in explaining confidence-accuracy dissociations. Therefore, we tested whether confidence-accuracy dissociations induced by stimulus energy manipulations emerge naturally in convolutional neural networks (CNNs). We found that, across three different energy manipulations, CNNs produced confidence-accuracy dissociations similar to those found in humans. This effect was present for a range of CNN architectures from shallow 4-layer networks to very deep ones, such as VGG-19 and ResNet -50 pretrained on ImageNet. Further, we traced back the reason for the confidence-accuracy dissociations in all CNNs to the same signal-and-variance increase that has been proposed for humans: higher stimulus energy increased the separation and variance of the CNNs’ internal representations leading to higher confidence even for matched accuracy. These findings cast doubt on the necessity of the positive evidence heuristic to explain human confidence and establish CNNs as promising models for adjudicating between low-level, stimulus-driven and high-level, cognitive explanations of human behavior.

## Introduction

Humans have the metacognitive ability to express confidence in their decisions (Koriat, 2006; Metcalfe & Shimamura, 1994). Although confidence is generally reliable in tracking one’s performance (Mamassian, 2016), several kinds of stimulus manipulations have been found to cause confidence to dissociate from accuracy (Boldt et al., 2017, 2019; de Gardelle & Mamassian, 2015; Desender et al., 2018; Herce Castañón et al., 2019; Koizumi et al., 2015; Samaha et al., 2016; Spence et al., 2016, 2018; Zylberberg et al., 2014, 2016).

A particular type of confidence-accuracy dissociations has been induced by stimulus manipulations referred to as “energy manipulations” (Gao et al., 2023; Zylberberg et al., 2012). Specifically, high energy stimuli can be created by simultaneously increasing stimulus features that aid recognition (e.g., stimulus contrast, motion coherence, etc.) and stimulus features that impede recognition (e.g., variability among several stimuli, strength of disconfirming evidence, etc.). For instance, Herce Castañón et al. (2019) manipulated stimulus energy by changing the contrast of an array of Gabor patches along with the variance of the orientations across the array, with observers having to decide on the overall orientation. Similarly, Koizumi et al. (2015) manipulated stimulus energy by increasing the contrast of both a target and a non-target superimposed grating, with observers having to pick the grating with higher contrast. These and similar types of energy manipulations are known to lead to confidence-accuracy dissociations, such that high stimulus energy leads to higher confidence in spite of accuracy being matched across energy levels (de Gardelle & Mamassian, 2015; Herce Castañón et al., 2019; Koizumi et al., 2015; Samaha et al., 2016; Spence et al., 2016; Zylberberg et al., 2014, 2016). For simplicity, in the rest of the paper we refer to stimulus features that aid recognition as “contrast” and stimulus features that impede recognition as “variability” because these terms describe well the majority of designs we examine in this study.

Explanations of these types of dissociations typically invoke “high-level” mechanisms. The most popular explanation is the positive evidence heuristic which assumes that confidence neglects evidence that disconfirms the observer’s choice (Maniscalco et al., 2016; Odegaard et al., 2018; Peters et al., 2017; Samaha et al., 2016; Zylberberg et al., 2012). The positive evidence heuristic predicts higher confidence for high-energy stimuli because high-energy stimuli lead to more extreme positive (as well as negative) evidence and confidence ignores negative evidence. Alternatively, these dissociations have been explained by assuming that humans infer decisions from an incorrect internal model of the task or themselves, which can result in suboptimal confidence. Particularly, observers’ internal models have been proposed to be either insensitive to stimulus variance (Zylberberg et al., 2014) or “blind” to noise arising from one’s own cognitive processes (Herce Castañón et al., 2019). In sum, these findings have been taken as evidence for confidence computations being based on a high-level inference process or heuristics.

On the other hand, a simpler “low-level” explanation for these effects posits that energy manipulations lead to changes in low-level perceptual representations, naturally leading to higher confidence (Fetsch et al., 2014; Morales et al., 2015; Rahnev et al., 2011, 2012, 2013; Zylberberg et al., 2016). Mechanistically, an increase in stimulus energy could lead to greater separation between evidence distributions a well as higher overall variability in the observed evidence – which we call the signal-and-variance-increase hypothesis. Consequently, a larger proportion of this distribution is shifted towards extreme values, thus increasing overall confidence (Gao et al., 2023). Critically, this explanation does not rely on higher-order mechanisms for producing the confidence-accuracy dissociations observed with energy manipulations.

Despite their importance for understanding the processes that give rise to confidence, it has been challenging to adjudicate between the “high-” and “low-” level explanations. The reason is that both explanations can fit the data, but there is no direct way of testing the assumptions inherent in each explanation.

Here, we use convolutional neural networks (CNNs) to distinguish between the “high-“ and “low-“ level explanations of the confidence-accuracy dissociations induced by stimulus energy manipulations. CNNs lack the built-in “high-level” cognitive mechanisms that are assumed to be responsible for confidence-accuracy dissociations. Therefore, if cognitive processes such as the positive evidence heuristic (Koizumi et al., 2015; Maniscalco et al., 2016; Peters et al., 2017; Samaha et al., 2016; Webb et al., 2023) or inference from suboptimal internal models (Herce Castañón et al., 2019; Zylberberg et al., 2014) are indeed necessary for these dissociations, these networks should fail to mimic human behavior. On the other hand, if confidence-accuracy dissociations arise from a low-level signal-and-variance increase based on inherent stimulus or task characteristics (Fetsch et al., 2014; Morales et al., 2015; Rahnev et al., 2011, 2012, 2013; Zylberberg et al., 2016), we can expect neural network models to reproduce human behavior. An additional advantage of CNNs is that, unlike humans, we can directly probe the network’s internal representations and understand the mechanisms underlying their behavior.

In this study, we tested whether three CNN architectures (a custom 4-layer CNN, VGG-19, and ResNet-50) produce human-like confidence accuracy dissociations across three types of energy manipulations. To anticipate, we found that all networks, like humans, expressed higher confidence for higher stimulus energy levels in spite of accuracies being matched. In addition, they reproduced another signature of confidence that is popularly regarded as evidence for the positive evidence bias (Maniscalco et al., 2016; Webb et al., 2023). Further, we show that the confidence increase in the CNNs was due to an increase in separability and variance of evidence distributions, which is essentially the signal-and-variance increase hypothesis that has been proposed for humans too (Fetsch et al., 2014; Gao et al., 2023; Morales et al., 2015; Rahnev et al., 2011, 2012, 2013; Zylberberg et al., 2016). These results demonstrate that CNNs exhibit human-like dissociations between confidence and accuracy, suggesting that higher-order mechanisms are unnecessary to explain the stimulus energy-induced effects on confidence. Importantly, these observations highlight the common mechanisms underlying the behavior of humans and artificial systems, suggesting that these confidence-accuracy dissociations may be driven by external features of the environment, without the need for specialized internal cognitive mechanisms.

## Results

We tested three CNN architectures (a custom 4-layer CNN, VGG-19, and ResNet-50) on three experiments involving two-choice discrimination judgements about the orientation of stimuli (**Figure 1**). The deep CNNs – VGG-19 and ResNet-50 – were pretrained on the ImageNet dataset and fine-tuned to perform these tasks. Experiments 1 and 2 have previously been shown to generate confidence-accuracy dissociations in humans (Herce Castañón et al., 2019; Koizumi et al., 2015), while Experiment 3 involved a novel task paradigm that has previously not been tested on either humans or neural networks. In each experiment, the energy of the stimulus was manipulated by simultaneously varying two independent features: the contrast of the stimulus and either its variability (in Experiments 1 and 3) or the contrast of the stimulus for the wrong choice (in Experiment 2). For all experiments, we trained 25 instances of each network architecture on 10,000 training images over a wide range of stimulus parameters and tested them on 1000 images from each energy condition. The stimulus parameters were chosen such that increasing energy levels resulted in the same average performance level of ∼70% across the 25 instances of each network architecture.

**Figure 1.**
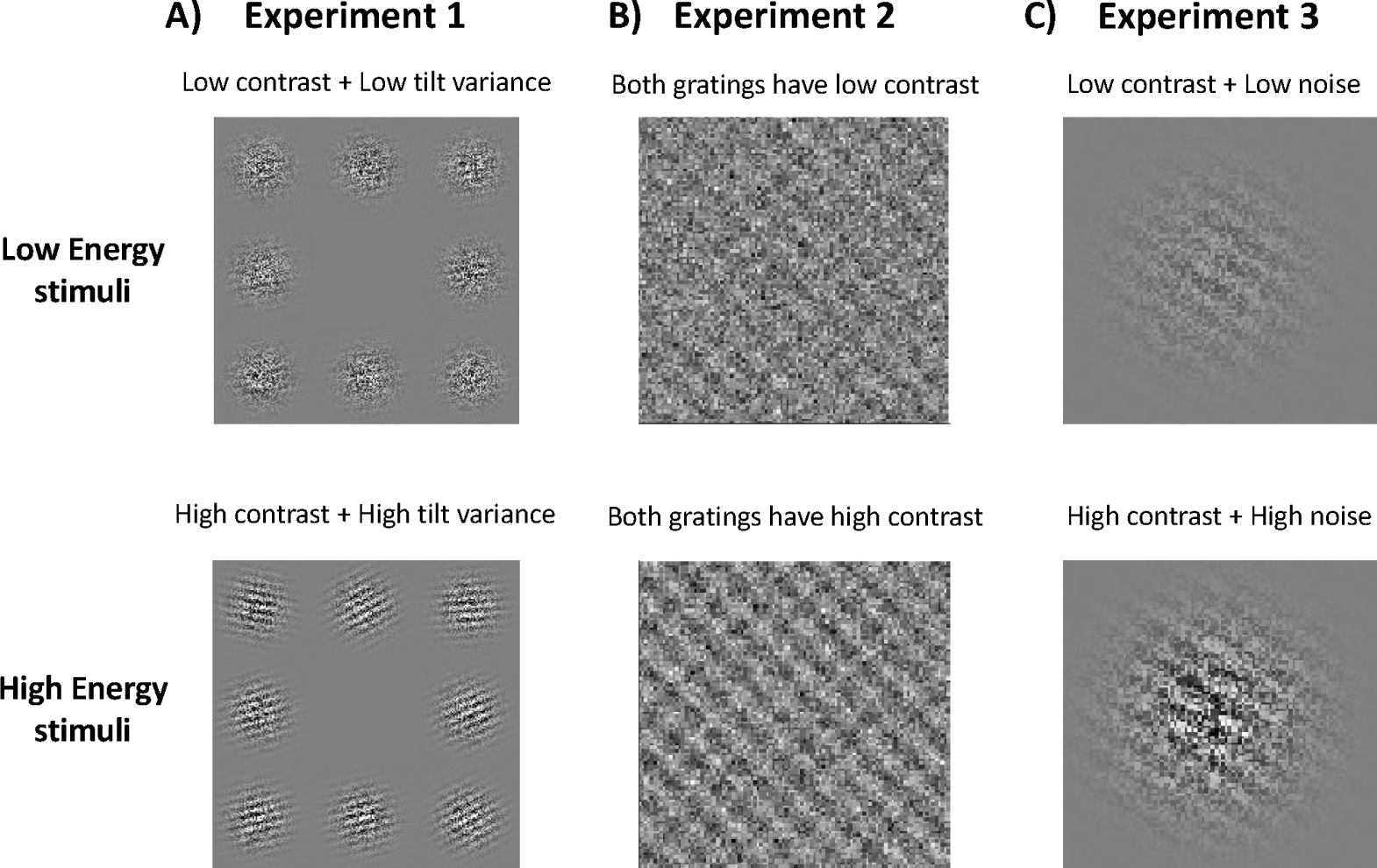
Energy manipulations. In all three experiments, the task involved two-choice discrimination between clockwise and counterclockwise oriented stimulus configurations. The upper and lower panels show examples of low and high energy stimuli respectively for each experiment. In all three examples, the correct choice is “counterclockwise”. (A) Task used by Herce Castañón et al. (2019). The stimulus consisted of an array of eight noisy Gabor patches with the task involving judgements of mean orientation relative to horizontal. Energy manipulations involved jointly changing the contrast of Gabors as well the variability of orientations across the array. (B) Task used by Koizumi et al. (2015). The stimulus consisted of two superimposed sinusoidal gratings overlaid by a noise mask. The task was to determine the orientation of the grating with the higher contrast (dominant grating). Increases in energy involved jointly increasing the contrast of the dominant and the non-dominant gratings. (C) The stimulus was a single Gabor patch overlaid with noise and the task was to determine its orientation. Energy was manipulated by jointly changing the contrast and noise level in the patch.

### CNNs exhibit robust confidence-accuracy dissociations

We computed the average accuracy and confidence of the 25 network instances for each of the three CNN architectures. First, we confirmed that we successfully matched model accuracies across the three energy conditions (**Figure 2**). Indeed, one-way repeated-measures ANOVAs on model accuracy with energy as factor showed that there were no significant differences in accuracy across the three energy conditions for all experiments and across all three CNN architectures: 4-layer CNN (Experiment 1: F(2,24) = .07, p = .93; Experiment 2: F(2,24) = 2.80, p = .07; Experiment 3: F(2,24) = .89, p = .42), VGG-19 (Experiment 1: F(2,24) = 2.08, p = .14; Experiment 2: F(2,24) = 1.98, p = .15; Experiment 3: F(2,24) = .76, p = .47), and ResNet-50 (Experiment 1: F(2,24) = .60, p = .56; Experiment 2: F(2,24) = .67, p = .52; Experiment 3: F(2,24) = .48, p = .62).

**Figure 2.**
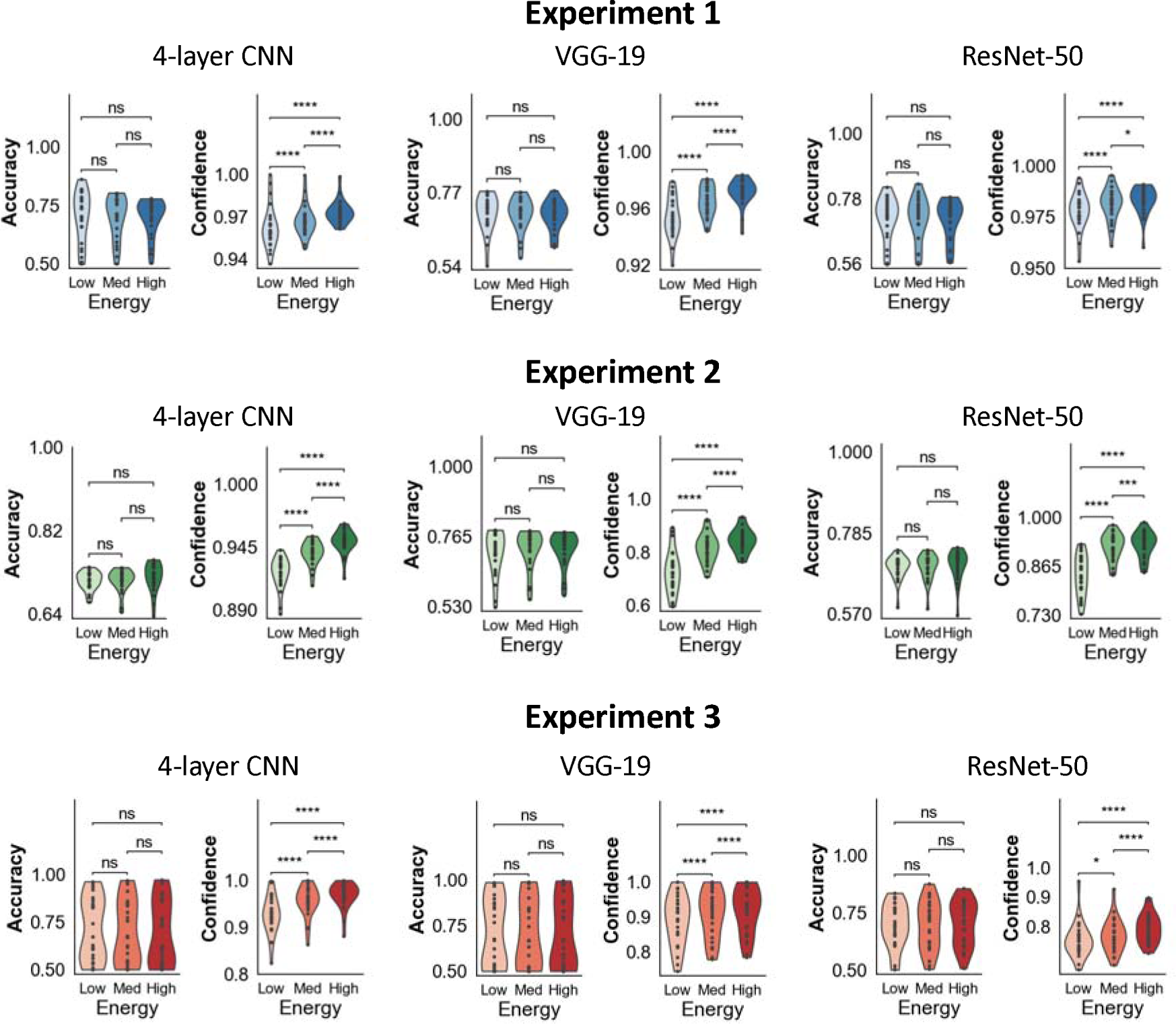
Confidence-accuracy dissociations in CNNs. For all experiments and networks (custom 4-layer CNN, VGG-19 and ResNet-50), accuracy was matched across energy conditions but confidence significantly increases with energy levels. Dots refer to individual network instances. The violin plots show the kernel density estimates of the data distribution. *p<0.05; **p<0.01; ***p<0.001; ****p<0.0001; n.s., not significant.

On the other hand, we found that increasing stimulus energy led to significantly higher confidence (**Figure 2**). Indeed, one-way repeated-measures ANOVAs showed highly significant differences in model confidence across the three energy conditions for all experiments in each of the three network architectures: 4-layer CNN (Experiment 1: F(2,24) = 36.27, p < .0001; Experiment 2: F(2,24) = 189.77, p < .0001; Experiment 3: F(2,24) = 51.82, p < .0001), VGG-19 (Experiment 1: F(2,24) = 85.60, p < .0001; Experiment 2: F(2,24) = 155.53, p < .0001; Experiment 3: F(2,24) = 39.15 p < .0001), and ResNet-50 (Experiment 1: F(2,24) = 20.75, p < .0001; Experiment 2: F(2,24) = 175.36, p < .0001; Experiment 3: F(2,24) = 19.85, p < .00001). Further, pairwise comparisons between the low and high energy levels showed highly significant increases in confidence for the high energy condition for all networks and all three experiments (all p’s < 0.0001; **Figure 2**). These results mimic the findings from human behavior where increasing the energy of the stimulus leads to increases in confidence, in spite of accuracies being matched across conditions (Boldt et al., 2017, 2019; de Gardelle & Mamassian, 2015; Desender et al., 2018; Herce Castañón et al., 2019; Koizumi et al., 2015; Samaha et al., 2016; Spence et al., 2016, 2018; Zylberberg et al., 2014, 2016). These findings cast doubt on the necessity of high-level explanations of confidence involving the positive evidence and noise-blindness hypotheses and are in-line with the predictions of the low-level signal-and-variance-increase hypothesis (Fetsch et al., 2014; Gao et al., 2023; Morales et al., 2015; Rahnev et al., 2011, 2012, 2013; Zylberberg et al., 2016). We note that in most cases, the CNNs exhibit average confidence levels greater than 0.9 in spite of mean accuracy being around 70%. These findings are in line with observations that neural networks, particularly CNNs, often exhibit overconfidence in their responses (Guo et al., 2017; Minderer et al., 2021).

### Mechanism behind the confidence-accuracy dissociation in CNNs

While findings of confidence-accuracy dissociations in CNNs support the signal-and-variance-increase hypothesis (Fetsch et al., 2014; Gao et al., 2023; Morales et al., 2015; Rahnev et al., 2011, 2013; Rahnev, Maniscalco, et al., 2012; Zylberberg et al., 2016),a more direct test of this hypothesis can be derived from examining how changes in stimulus energy affect the CNNs’ internal distributions. Specifically, the signal-and-variance-increase hypothesis predicts that increasing stimulus energy leads to greater separation of evidence between the two stimulus categories as well as an increase in the variance of evidence. As a result of these changes, the evidence distributions shift towards more extreme values, leading to higher confidence overall. Therefore, we probed the CNNs’ internal evidence representations to examine whether changes in the network’s internal representations are consistent with this hypothesis.

We accessed the CNNs’ internal representations by aggregating the activations generated in the networks’ output layer in response to images for each stimulus category separately for each energy condition. We then plotted these activations for each stimulus category (*S*_1_ for counterclockwise stimuli and *S*_2_ for clockwise stimuli) and observed how the characteristics of these distributions vary across energy levels. We quantified the separation between the two stimulus categories as the distance between their means (λ*S*_2_ – λ*S*_1_; where *ν_S__i_* refers to the mean of the evidence distribution for stimulus category *i*) and the spread of distributions as the average standard deviation (SD) of the two evidence distributions. The separation between distributions and the average SD was computed separately for each of the 25 network instances.

We found that increasing energy levels led to larger separation between the *S*_1_ and *S*_2_ evidence distributions as well as an increase in the variance of these distributions (**Figure 3**). Indeed, one-way repeated measures ANOVAs showed highly significant differences in the separation between the two distributions between the three energy conditions for all experiments and networks: 4-layer CNN (Experiment 1: F(2,24) = 320.83, p < .0001; Experiment 2: F(2,24) = 342.21, p < .0001; Experiment 3: F(2,24) = 92.32, p < .0001), VGG-19 (Experiment 1: F(2,24) = 169.78 p < .0001; Experiment 2: F(2,24) = 177.81, p < .0001; Experiment 3: F(2,24) = 160.13, p < .0001), and ResNet-50 (Experiment 1: F(2,24) = 444.34, p < .0001; Experiment 2: F(2,24) = 121.60, p < .0001; Experiment 3: F(2,24) = 54.37, p < .00001). Further, pairwise comparisons showed a significant increase in the separation between the distributions for the two stimulus categories from the low-to high-energy conditions for all experiments and networks (all p’s < 0.0001).

**Figure 3.**
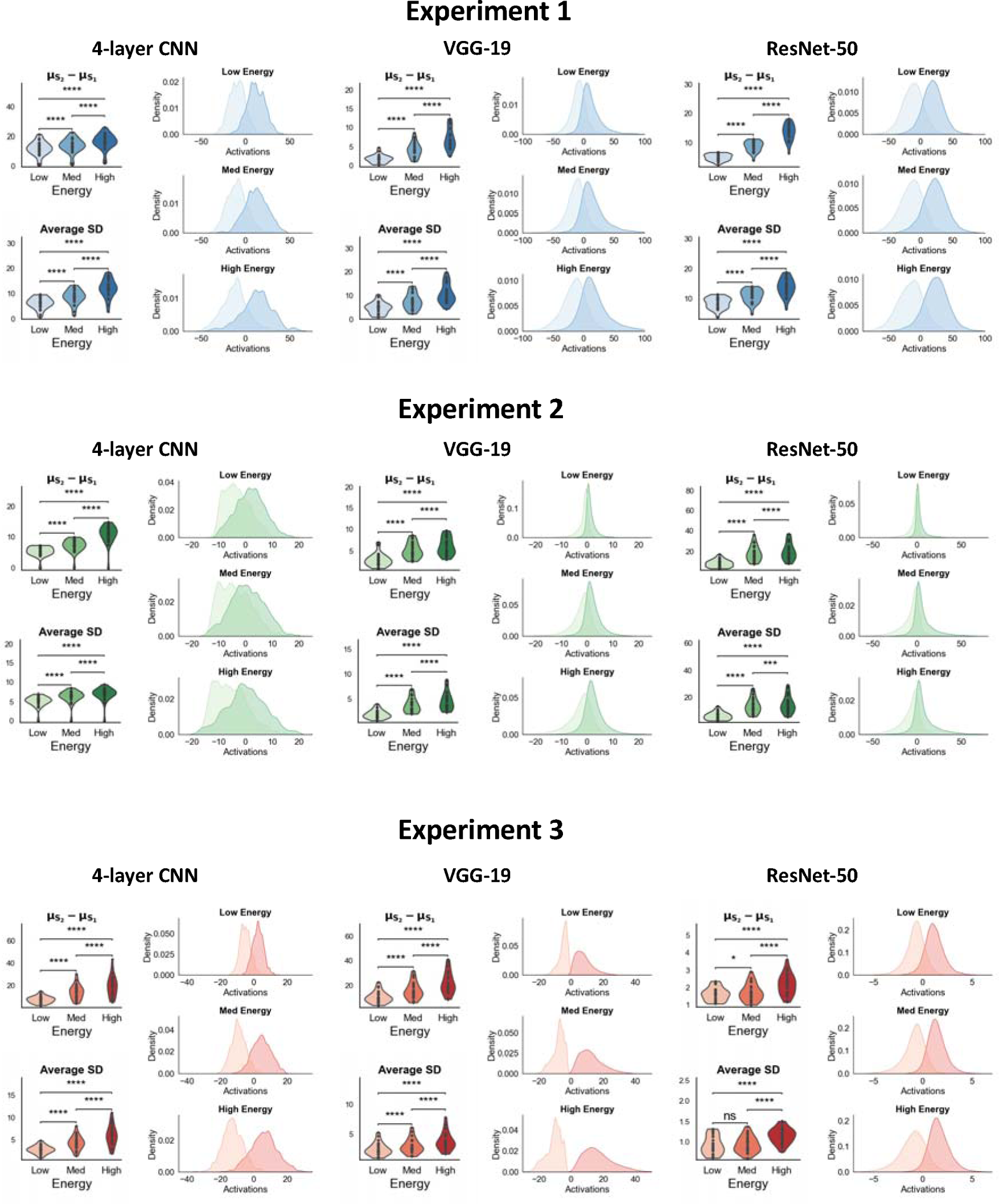
Energy manipulations increase both the separability and variance of the networks’ internal activations. For all three experiments, the separability between the distributions of internal evidence for the two stimulus categories, as well as the variance of the evidence distributions, increased with energy levels For each network, the figure shows the distance between the *S*_1_ and *S*_2_ evidence distributions and their standard deviations (SD) across the 25 model instances. The kernel density plots show the distribution of activations aggregated across all 25 network instances. *p<0.05; **p<0.01; ***p<0.001; ****p<0.0001; n.s., not significant.

Parallel to the results on separability, we found that higher stimulus energy also led to increases in the variance of the internal evidence distributions (**Figure 3**). Indeed, one-way ANOVAs on the average SD of evidence distributions also yielded highly significant differences between the three energy conditions for all experiments and networks: 4-layer CNN (Experiment 1: F(2,24) = 408.38, p < .0001; Experiment 2: F(2,24) = 284.13, p < .0001; Experiment 3: F(2,24) = 120.15, p < .0001), VGG-19 (Experiment 1: F(2,24) = 140.25, p < .0001; Experiment 2: F(2,24) = 108.84, p < .0001; Experiment 3: F(2,24) = 177.82 p < .0001), and ResNet-50 (Experiment 1: F(2,24) = 229.60, p < .0001; Experiment 2: F(2,24) = 108.84.38, p < .0001; Experiment 3: F(2,24) = 69.99, p < .00001). Further, pairwise comparisons across low and high energy conditions revealed significant increases in the average SD of activations for all networks and across all three experiments (all p’s < 0.0001).

The concurrent increase in separation between the two stimulus categories and the variability of evidence ensures that the network’s overall stimulus sensitivity remains constant between the three energy conditions. Specifically, any improvement in sensitivity yielded by the increased separation between the evidence distributions for the two categories is counteracted by the evidence itself becoming more variable. Nevertheless, confidence differences still emerge between conditions because the higher variance and separation between these distributions results in larger proportions of evidence being pushed towards extreme values that get assigned higher confidence. These findings show that the signal-and-variance-increase mechanism indeed underlies the confidence-accuracy dissociations produced by the CNNs.

### Different stimulus features selectively influence the separability and spread of internal evidence distributions

While changes in characteristics of the stimulus distributions can explain the differences in confidence between energy conditions, it is still unclear why energy manipulations affect these characteristics. To address this question, we compared the networks’ internal representations during energy manipulations to the representations resulting from manipulations of contrast and variability (Experiments 1 and 3) or contrast of the correct vs. incorrect grating (Experiment 2). For conciseness, we refer to manipulations of both variability and the contrast of the incorrect grating as manipulations of variability. For these analyses, we only focus on the activations of the simplest network – the 4-layer CNN – although we expect these results to generalize across other network architectures. For each experiment, we investigated the network’s activations separately for manipulations of contrast and manipulations of variability.

We first examined the effects of contrast and variability manipulations on the separability of the internal evidence distributions. We found that while increasing stimulus contrast increased the separation between stimulus categories, increasing variability led to a decrease in their separation (**Figure 4A**). Indeed, there were significant mean differences in evidence separability for both manipulations of contrast (Experiment 1: F(2,24) = 233.44, p < .0001; Experiment 2: F(2,24) = 283.89, p < .0001; Experiment 3: F(2,24) = 239.62, p < .0001) and manipulations of variability (Experiment 1: F(2,24) = 57.75, p < .0001; Experiment 2: F(2,24) = 324.00, p < .0001; Experiment 3: F(2,24) = 763.58, p < .0001). Pairwise comparisons showed that contrast significantly increased separability between stimulus categories (all p’s < .0001, paired t-tests comparing the lowest and highest contrast levels), while variability significantly decreased this separation (all p’s < .0001, paired t-tests comparing the highest and lowest contrast levels). These results suggest that the increase in separability between evidence distributions observed during energy manipulations is primarily driven by changes in stimulus contrast.

**Figure 4.**
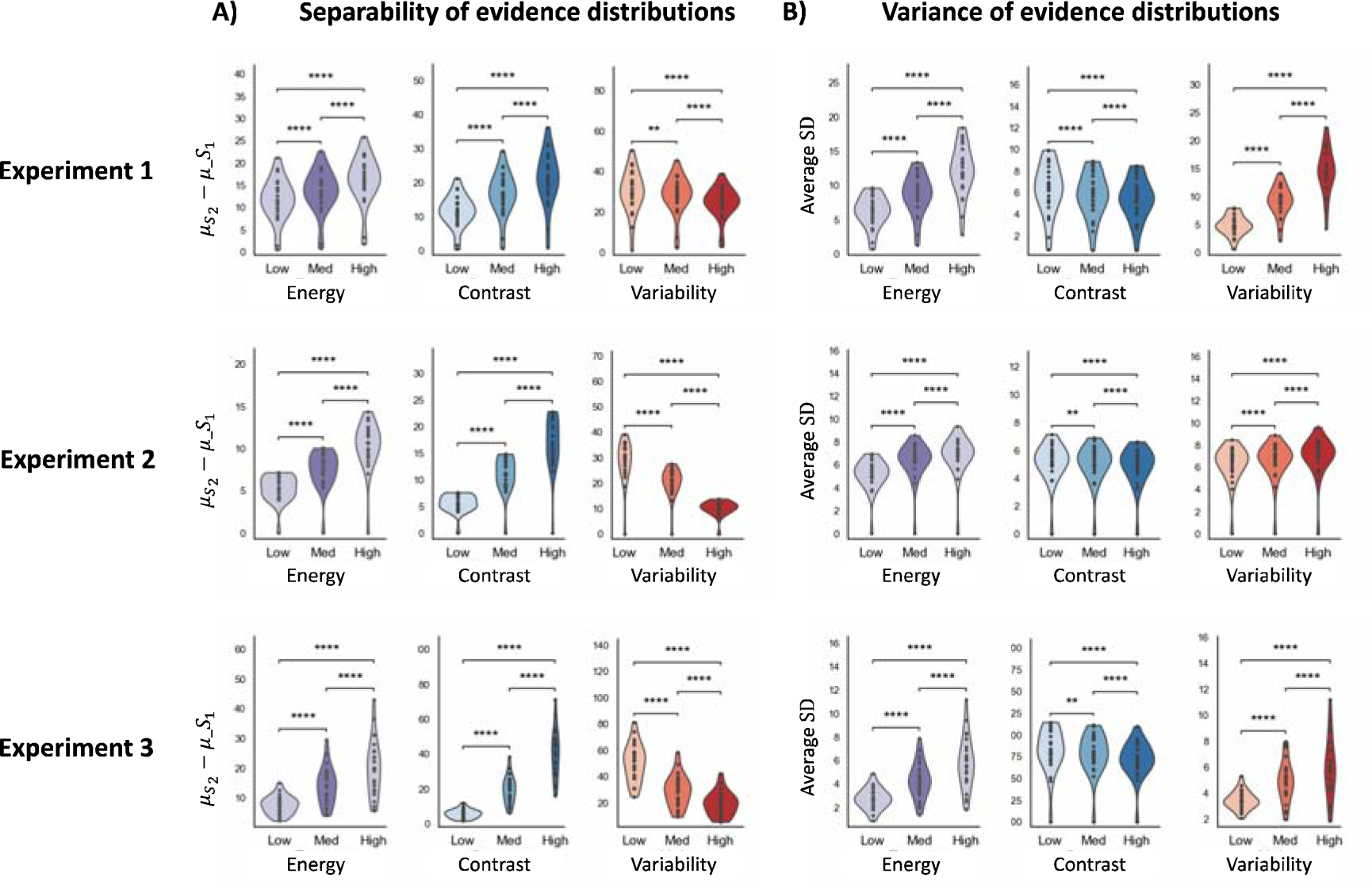
Changes in the separation and spread of internal evidence distributions induced by energy, contrast, and variability manipulations. The plots show A) the average distance between the mean activations for *S*_1_ and *S*_2_ stimuli (left) and B) the average standard deviation (SD) of activations (right) in the final layer of the shallow CNNs for Experiments 1-3 in response to changes in stimulus energy, contrast and variability. Note that the energy results in both panels are equivalent to the 4-layer CNN results from Figure 3. For all experiments, increasing energy and contrast levels increases the separation between the two stimulus categories, while increasing variability decreases the separability between the two stimulus categories. On the other hand, increasing stimulus energy and variability increases the spread of evidence distributions, while increasing contrast decreases the spread of evidence. These results suggest contrast and noise changes selectively drive changes in separation and variance of evidence distributions respectively. The violin plots show the kernel density estimates of the data distribution. *p<0.05; **p<0.01; ***p<0.001; ****p<0.0001; n.s., not significant.

We then examined the effects of contrast and variability manipulations on the spread of the internal evidence distributions. We found that increasing stimulus contrast decreased the variance of evidence distributions, while increasing variability increased their variance. Indeed, there were significant differences in the spread of evidence distribution for both manipulations of contrast (Experiment 1: F(2,24) = 56.03, p < .0001; Experiment 2: F(2,24) = 21.71, p < .0001; Experiment 3: F(2,24) = 113.61, p < .0001) and manipulations of variability (Experiment 1: F(2,24) = 388.45, p < .0001; Experiment 2: F(2,24) = 48.16, p < .0001; Experiment 3: F(2,24) = 53.28, p < .0001). Pairwise comparisons showed that variability significantly increased the spread of distributions (paired t-tests comparing the highest and lowest contrast levels; all p-values < .0001) while contrast significantly decreased their spread (paired t-tests comparing the highest and lowest contrast levels; all p-values < .0001). These results suggest that the increase in variability of evidence observed during energy manipulations is primarily driven by changes in stimulus variability (**Figure 4; right**). Overall, these results demonstrate that manipulations of stimulus contrast and variability have opposite effects on the separability and spread of internal activations, such that contrast manipulations have a larger effect on separability and variability manipulations have a larger effect on spread. Thus, combining both manipulations in a single “energy” manipulation leads to both increased separability and spread of the internal evidence distributions.

### CNNs can reproduce dissociations between type-1 and type-2 sensitivity typically regarded as evidence for the positive evidence mechanism

So far, our results have shown that CNNs, in spite of lacking the positive evidence mechanism, can produce human-like confidence-accuracy dissociations that have typically been attributed to the positive evidence heuristic. Another feature of confidence that has been attributed to the positive evidence mechanism is the observation that under certain conditions, an observer’s type-1 and type-2 sensitivities are found to dissociate from each other (Maniscalco et al., 2016; Webb et al., 2023). Here, type-1 sensitivity (d’) refers to the amount of information available for the primary stimulus judgement, whereas type-2 sensitivity (meta-d’) reflects the amount of information underlying confidence judgements. Typically, an increase in type-1 sensitivity translates into a proportional increase in type-2 sensitivity. However, Maniscalco et al. (2016) found that under certain task paradigms, these two measures can dissociate from each other. More specifically, the paradigm consists of a two-choice discrimination task, where the contrast of one stimulus category (*S*_1_) is held constant while the contrast of the other stimulus is allowed to vary over discrete levels (*S*_2_). Under these conditions, meta-d’ decreases with d’ for trials where the observer responds “*S*_1_,” but meta-d’ increases with d’ for trials where the observer responds “*S*_2_.” Importantly, this effect was explained by a model incorporating the positive evidence mechanism, whereas a competing model that assumed equal weights for positive and negative evidence failed to account for this behavior (Maniscalco et al., 2016). These findings are typically regarded as evidence for the existence of the positive evidence mechanism for confidence.

To further test the necessity of the positive evidence mechanism in explaining confidence, we assessed whether CNNs lacking this mechanism can also reproduce the dissociation observed in Maniscalco et al. (2016). We simulated the above task paradigm for our previously trained 4-layer CNNs across the three experiments (as done by Webb et al., 2023) and found that for Experiments 1 and 3, the networks were indeed able to reproduce a clear dissociation between meta-d’ and d’ (**Figure 5**). Specifically, meta-d’ increases with d’ for trials with “*S*_1_” responses (where the contrast of *S*_2_ varies across trials), but meta-d’ decreases with d’ for trials with “*S*_1_” responses (the contrast of “*S*_1_ remaining fixed across trials), producing the distinct cross-over signature shown by Maniscalco et al. (2016). However, for Experiment 2, while meta-d’ increased steeply with d’ for trials where the observer responds “*S*_2_,” meta-d’ also showed a slight increase with d’ for trials where the observer responds “*S*_1_.” These results suggest that unlike the confidence-accuracy dissociations, the meta-d’/d’ dissociations may be more sensitive to the specific characteristics of the stimuli. Overall, our findings demonstrate that for at least for some stimulus manipulations, low-level mechanisms are sufficient to explain effects that have typically been taken as evidence for a positive evidence mechanism.

**Figure 5.**
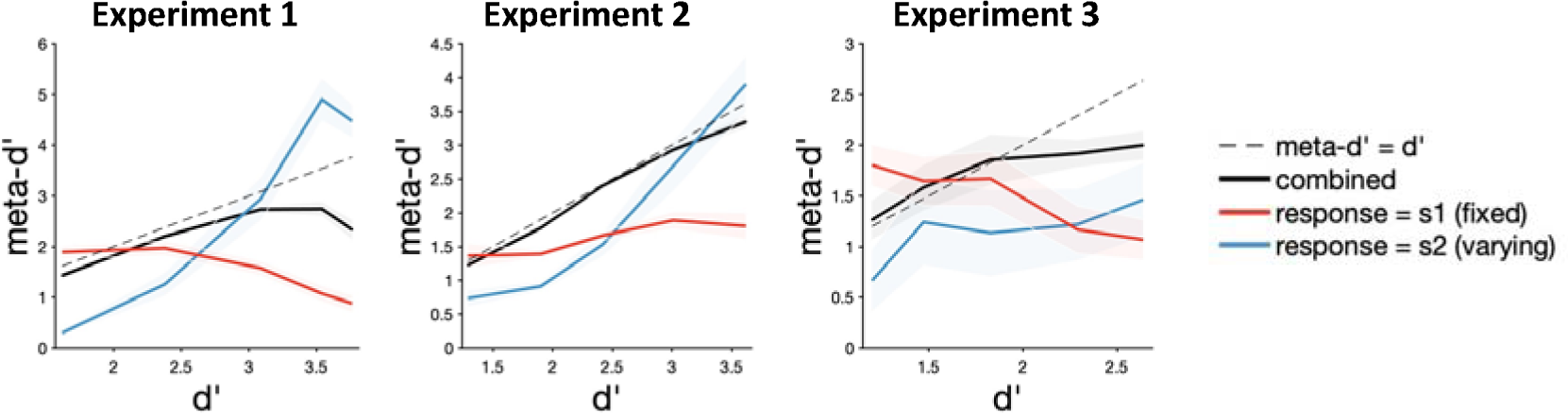
Dissociations between meta-d’ and d’ in 4-layer CNNs. We tested the 4-layer CNNs on the task paradigm from Webb et al. (2023) which demonstrated a dissociation between d’ and meta-d’ under certain conditions. The contrast of one stimulus category (*S*_1_) remains fixed while the contrast of the other stimulus is increased in discrete steps (*S*_2_) Meta-d’ increases with d’ as expected for trials in which the observer responds “*S*_2_”, but meta-d’ decreases with d’ while for trials where the observer responds “*S*_1_”. This behavioral effect is predicted by a model incorporating the positive-evidence bias. Here, we simulated this task paradigm for Experiments 1-3. The responses generated by our 4-layer CNNs show that these networks can indeed generate the meta-d’-d’ dissociations observed in humans for Experiments 1 and 3. The network fails to reproduce this behavior for Experiment 2, suggesting that these dissociations may depend on the specifics of the stimuli used for the tasks.

### Confidence accuracy dissociations in CNNs generalize across stimulus paradigms but do not always mimic human behavior

Our results demonstrate that CNNs can produce human-like confidence-accuracy dissociations where confidence increases with increasing stimulus energy levels. However, when using color stimuli, energy manipulations have been found to *decrease* confidence while accuracy remains matched across conditions (Boldt et al., 2017; de Gardelle & Mamassian, 2015; Desender et al., 2018; Spence et al., 2016, 2018). These findings have been explained by assuming a mechanism of “robust averaging” where highly atypical stimuli are down-weighted in the final decision (Boldt et al., 2024; De Gardelle & Summerfield, 2011).

Our previous findings establish that low-level changes in perceptual representations can explain human behavior that has typically been attributed to high-level cognitive mechanisms. Here, we sought to further test whether the “robust averaging” mechanism can also be realized through low-level mechanisms.

Following the same procedure as done previously, we tested our CNNs on the task from Boldt et al. (2017) where subjects identified whether the mean color across an array of eight colored patches was closer to red or blue (**Figure 6A**). Energy manipulations involved jointly increasing the intensity of the color (“blueness” or “redness” of stimuli) and the variance of color across the eight patches. The stimulus parameters were chosen such that we obtained matched average accuracy levels of ∼70% across the three energy levels. A one-way repeated measures ANOVA showed no significant mean differences in accuracy between the three energy conditions for all three networks – 4-layer CNN (F(2,24) = .78, p = .47), VGG-19 (F(2,24) = .31, p = .74) and ResNet-50 (F(2,24) = .15, p = .86).

**Figure 6.**
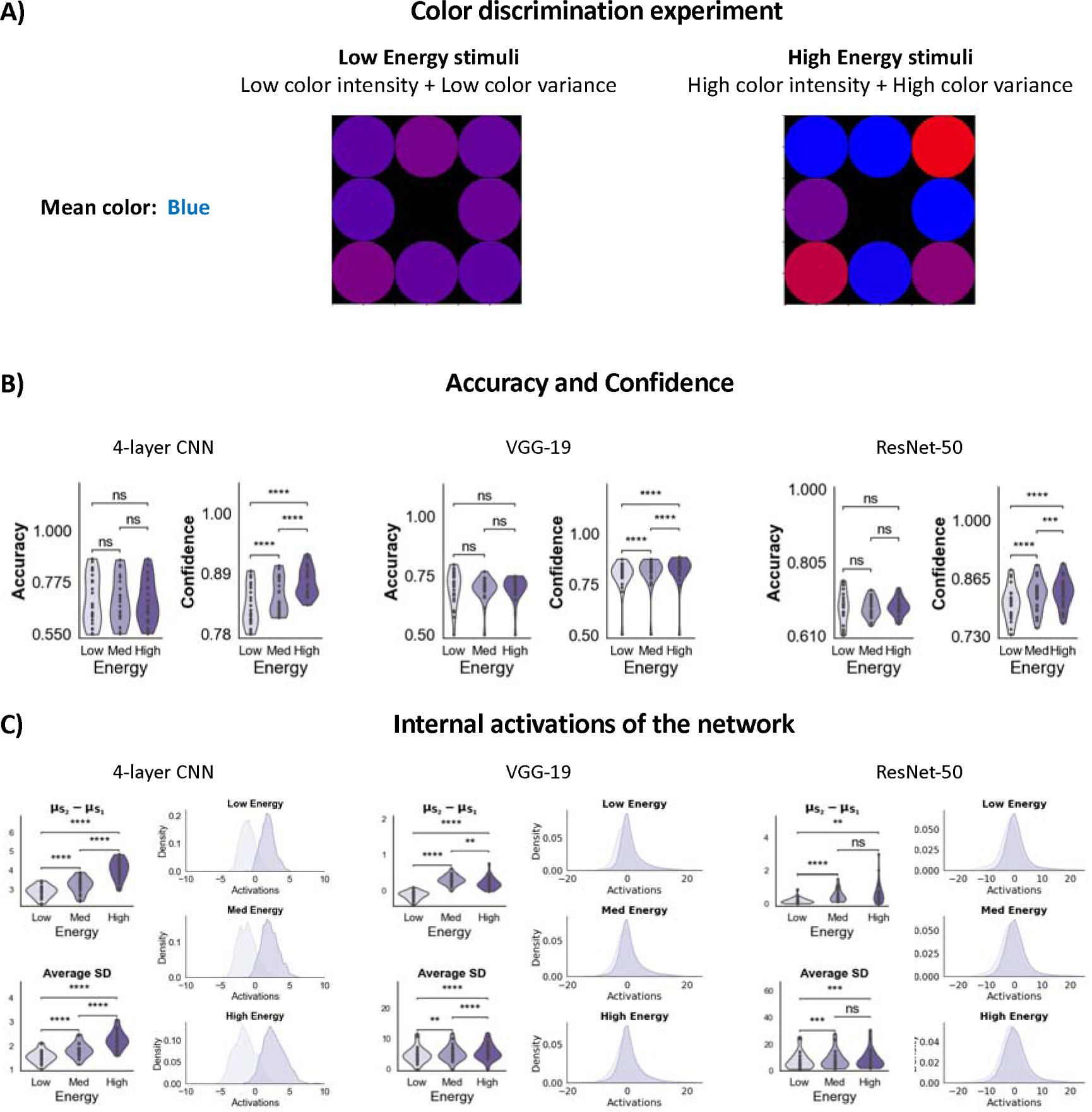
Confidence-accuracy dissociations in a color discrimination task. A) The stimulus consisted of an array of eight colored circles. The task was to determine whether the mean color across the eight patches was more blue or red. In this example, the mean color is more blue than red. Energy manipulations involved joint changes to the intensity of color (the amount of “blueness” or “redness” of the patches as well the variance in color across the array. (B) The separability between the stimulus categories as well as the variance of the evidence distributions increased with energy levels for all three networks. The panels on the top-left for each network show the average distance between the *S*_1_ and *S*_2_ evidence distributions across the 25 model instances. The panels on the bottom-left show the average standard deviation (SD) across the two distributions across all model instances. The panels on the right show the distribution of activations aggregated across all 25 network instances. *p<0.05; 250 **p<0.01; ***p<0.001, ****p<0.0001; n.s., not significant.

However, the ANOVA revealed highly significant increases in confidence between the three energy levels for all three networks (4-layer CNN: F(2,24) = 152.50, p < .0001; VGG-19: F(2,24) = 57.72, p < .0001; and ResNet-50: F(2,24) = 68.71, p < .0001; pairwise comparisons between low and high energy levels: all p’s < 0.0001; **Figure 6B**). Further, these behavioral effects were associated with increases in both the separability and variance of evidence distributions (all pairwise comparisons between low and high energy levels < .002; **Figure 6C**), in line with the signal-and-variance-increase effect. While these results are consistent with findings from Experiments 1-3, they fail to replicate human behavior suggesting that low-level explanations cannot account for all types of confidence-accuracy dissociations, particularly, the ones attributed to a process of “robust averaging”. Future studies must investigate whether incorporating the robust averaging mechanism can restore human-like confidence behavior in CNNs. Nevertheless, these results while establishing the generalizability of the mechanisms underlying confidence-accuracy dissociations in CNNs also reveal how stimulus-specific interactions can constrain the similarities between humans and CNNs. Understanding the conditions that separate the behavior of humans and CNNs can guide future work on building metacognitive systems for AI.

## Discussion

We found that convolutional neural networks (CNNs) robustly produce human-like confidence-accuracy dissociations in response to stimulus energy manipulations. In humans, these dissociations have been taken as evidence for high-level, cognitive mechanisms such as the positive evidence heuristic (Koizumi et al., 2015; Odegaard et al., 2018; Samaha et al., 2016)and noise blindness (Herce Castañón et al., 2019; Zylberberg et al., 2014). Since CNNs lack such built-in cognitive mechanisms, their ability to mimic human confidence behavior implies that these popular theories are unnecessary to explain energy-induced confidence-accuracy dissociations. Our findings support the alternative, signal-and-variance increase hypothesis which posits that such dissociations naturally emerge from low-level changes in perceptual representations. Indeed, we find that in CNNs, these dissociations are explained by their internal representations becoming more separable as well as variable with higher stimulus energy. These findings highlight how the behavior of both artificial and biological systems could be driven by common, stimulus-driven processes and demonstrate the usefulness of CNNs in distinguishing between low- and high-level explanations of behavior.

### Implications for the positive evidence bias in confidence

The positive evidence (PE) heuristic is one of the most popular proposals regarding the computations underling confidence (Koizumi et al., 2015; Maniscalco et al., 2016; Odegaard et al., 2018; Peters et al., 2017; Samaha et al., 2016; Webb et al., 2023; Zylberberg et al., 2012). Despite its popularity, however, findings from recent studies suggest that the positive evidence bias may not be necessary to explain confidence (Rausch et al., 2020; Shekhar & Rahnev, 2023).

Firstly, the previous studies that found support for the PE bias did not perform extensive model comparisons. Rather, the PE model was typically compared to a model which assumed that confidence is based on a balance of evidence between the two choice options (Maniscalco et al., 2016; Peters et al., 2017; Webb et al., 2023). Importantly, in this model stimulus manipulations were assumed to have no effect on the variance of the internal distributions of evidence. Further, the PE model has rarely been compared to other recently developed models of confidence in the literature (Bang et al., 2019; Boundy-Singer et al., 2023; Fleming & Daw, 2017; Guggenmos, 2022; Li & Ma, 2020; Mamassian & de Gardelle, 2021; Maniscalco & Lau, 2016; Rausch et al., 2020; Shekhar & Rahnev, 2021). When such model comparisons have been performed, the PE model usually ranks poorly relative to models that allow suboptimalities in confidence to manifest via other mechanisms such as metacognitive noise or the visibility heuristic (Rausch et al., 2020; Shekhar & Rahnev, 2023).

Secondly, behavioral evidence for the PE bias mainly rests on two observed patterns in confidence – increase in confidence with stimulus energy despite matched accuracies (Koizumi et al., 2015; Odegaard et al., 2018; Samaha et al., 2016) and the dissociation between d’ and meta-d’ under a certain stimulus paradigm (Maniscalco et al., 2016; Webb et al., 2023). However, these studies have not considered the existence of alternative mechanisms that can explain these behavioral effects. Here, we show that CNNs, which lack any confidence-specific mechanisms, can produce both these signatures via changes in their evidence representations. This result thus questions whether such behavioral findings can, by themselves, be taken as evidence for the positive evidence mechanism.

### Implications for other high-level theories of confidence

Findings of energy-induced confidence-accuracy dissociations have also been interpreted as evidence for other cognitive processes. For instance, Herce Castañón et al. (2019) argued that a range of suboptimal behaviors arise from noise blindness, such that observers neglect to account for the noise arising from their own cognitive computations when integrating across variable evidence samples. This noise blindness results in observers failing to adjust their responses to increasing levels of uncertainty. In addition to being over-confident for high-energy stimuli, Herce Castañón et al. report that observers neglect stimulus base rates in the high-energy condition and thus fail to appropriately shift their decision criterion in favor of the more frequent stimulus. However, the signal-and-variance-increase hypothesis is sufficient to explain both the suboptimal behaviors they report. According to the signal-and-variance-increase hypothesis, when the separation between the distributions and their variance is high, a criterion shift of same magnitude will have a smaller effect on choice probabilities compared to when the distributions have low separation and variance, thus appearing as if the observers have failed to shift their criteria appropriately. Indeed, simulations of criterion shifts under the signal-and-variance-increase hypothesis reproduced their reported effects (**Supplementary Figure 1**).

However, it must be noted that overconfidence and base-rate neglect both arise from the observers’ failure to scale their decision and confidence criteria in response to low-level perceptual changes. In that sense, this effect might reflect a blindness to changes in low-level representations at the decision stage. Nevertheless, this mechanism is distinct from the one proposed by Herce Castañón et al. (2019) because in their model the blindness is towards noise arising from the observers’ own internal cognitive processes, rather than towards noise arising from stimulus-driven changes in internal representations.

### Evidence for the signal-and-variance-increase hypothesis

The notion that confidence can be influenced by low-level changes in internal evidence representations is not new. Stimulating lower (Rahnev et al., 2012, 2013) and mid-level visual areas (Fetsch et al., 2014) have been found to affect confidence, independent of changes in accuracy. These effects were captured well by models where stimulation increased the variance of internal representations. Other task manipulations involving attention (Morales et al., 2015; Rahnev et al., 2011) and evidence volatility (Zylberberg et al., 2016) also produced similar dissociations in confidence that were accounted by changes in the trial-by-trial variance of sensory evidence. Our current findings corroborate these findings and extend them by adding energy manipulations to the list of factors that can produce confidence-accuracy dissociations via the signal-and-variance-increase mechanism.

### CNNs as models for understanding human vision

Several recent studies have argued that deep neural networks can provide meaningful insights into the goals and constraints that have shaped human perception (Blauch et al., 2021; Cao & Yamins, 2021; Dobs et al., 2022; Doerig et al., 2023; Gomez-Villa et al., 2019; Kell et al., 2018; Kell & McDermott, 2019; Richards et al., 2019; Wichmann & Geirhos, 2023). Indeed, our findings support this argument as they carry implications about the external constraints that may have shaped visual processing. For instance, if it is indeed true that the signal-and-variance-increase mechanism underlies the behavior of both humans and CNNs, this fact can reveal how common behaviors (due to common mechanisms of visual processing) can emerge in artificial and biological neural networks due to external, stimulus-defined constraints. Another critical advantage of CNNs is that they can allow us isolate bottom-up visual processes from the top-down mechanisms that serve cognition, since visual processing in standard CNNs occurs in the absence of specialized top-down influences. In addition, they can inform us about the possible stimulus and task representations that underlie such bottom-up stimulus processing (Green et al., 2024).

### Other types of confidence-accuracy dissociations

In the current study, we tested CNNs on a specific type of confidence-accuracy dissociation induced by energy manipulations. However, prior research has found that an abundance of factors can cause confidence to dissociate from accuracy. Some of these include motor preparation and execution (Fleming et al., 2015; Gajdos et al., 2019; Wokke et al., 2020), transcranial magnetic stimulation (Rahnev et al., 2012, 2016; Rounis et al., 2010; Shekhar & Rahnev, 2018; Xue et al., 2023), differences in pre-stimulus brain activity Bahdo, et al., 2012; Samaha et al., 2017), confidence history (Aguilar-Lleyda et al., 2021; Rahnev et al., 2015), attention (Rahnev et al., 2011; Wilimzig et al., 2008), arousal (Allen et al., 2016), and stimulus visibility (Rausch et al., 2018, 2020). However, in this study, we only chose to test energy-induced confidence accuracy dissociations as these manipulations can be readily applied to CNNs unlike those involving motor preparation, transcranial magnetic stimulation, attention, arousal, etc. Future studies can test the proposed mechanisms underlying other kinds of confidence-accuracy dissociations against suitable low-level explanations to gain insight into the true mechanisms of confidence.

Importantly, beyond the examples of high-energy stimuli leading to high confidence examined in this paper, there are two kinds of stimuli that break that rule. For these stimuli, energy manipulations lead to confidence that decreases with energy levels. Firstly, Spence et al. (2016, 2018) observed this effect for random dot motion stimuli. It is possible to explain this effect by assuming that increasing the variance of motion direction may deliver high-level cues regarding task difficulty. In turn, subjects may use these difficulty cues to decrease their confidence. Since our CNNs do not work on dynamic, dot motion stimuli, we could not test them on these stimuli. Secondly, Boldt et al. (2017, 2019) and Desender et al. (2018) found a similar effect for arrays of colored dots. When we tested our CNNs on these color stimuli (**Figure 6**), we found that low-level mechanisms cannot account for these effects, and thus it is likely that these manipulations engage high-level cognitive mechanisms. Indeed, these color tasks have been proposed to trigger “robust averaging” where observers down-weight highly atypical evidence samples (De Gardelle & Summerfield, 2011). Since high-energy stimuli generate more extreme evidence, ignoring (or down-weighting) them leads to a lower overall estimate of evidence for confidence. Future studies should test whether incorporating high-level cues about task difficulty and the robust averaging mechanism into CNNs can indeed generate this effect.

### Conclusion

In this study, we demonstrate that CNNs can generate human-like confidence-accuracy dissociations in response to stimulus energy manipulations via changes in the variance and separability of their internal evidence distributions. These findings cast doubt on the necessity of invoking high-level, cognitive explanations for this phenomenon – particularly the popular assumption that confidence is derived from a positive-evidence heuristic. Our results highlight the necessity of disentangling low- and high-level explanations of behavior and establish CNNs as promising models for generating and testing hypotheses about the mechanisms underlying human behavior.

## Methods

### Stimuli and task

We tested several convolutional neural networks on three main experiments. The task paradigms for Experiments 1 and 2 were adapted from Herce Castañón et al. (2019) and Koizumi et al. (2015). Both of these papers found confidence-accuracy dissociations in humans where confidence was found to increase with stimulus energy levels. To test the generality of our findings, we also included a novel task paradigm as Experiment 3 that has not been previously tested on humans but nevertheless uses the same kind of energy manipulations.

In Experiment 1, the stimuli (90 x 90 pixels) consisted of an array of eight noisy, oriented Gabor patches. Each individual Gabor patch in the array spanned 30 x 30 pixels. The task was to decide whether the average tilt across the 8 patches was clockwise (CW) or counterclockwise (CCW) from the horizontal (**Figure 1A**). For each image, the average orientation of the Gabor patches across the eight patches was selected from a Gaussian distribution. The energy of the stimulus was manipulated across three levels by simultaneously varying two features of the array – the contrast of individual gratings and the variability of orientations across the gratings. While increasing the contrast of the gratings allowed better stimulus visibility and made the task easier, increasing the variability of orientations increased the uncertainty regarding the mean orientation across the patches, thus making the task harder.

In Experiment 2, the stimuli (100 x 100 pixels) consisted of two noisy, sinusoidal gratings (oriented either 45° CCW or CW to the vertical) superimposed on each other. The two gratings were always oriented orthogonally to each other and one of the gratings had a higher contrast (referred to as the dominant grating). The task was to determine whether the dominant grating was oriented CCW or CW to the vertical (**Figure 1B**). The energy of the stimulus was manipulated by simultaneously varying the contrast levels of the dominant and the non-dominant grating across three levels. Increasing the contrast of the dominant grating contributed positive evidence making the task easier, while increasing the contrast of the non-dominant grating increased the level of contradictory or “negative evidence” making the task harder.

In Experiment 3, the stimulus consisted of a single noisy Gabor patch (100 x 100 pixels) oriented 45° either CCW or CW to the vertical (**Figure 1C**). The task was to identify the direction of tilt (CCW/CW). The energy of the stimulus was manipulated by varying both contrast and noise of the gratings. While increasing contrast makes the task easier, increasing noise degraded the stimulus, making the task harder.

### Generating the training and validation sets

For each experiment, we trained the networks on a set of 10,000 images. To allow the networks to learn generalizable representations of the stimuli, we generated images by sampling the stimulus parameters uniformly within a range. In Experiment 1, we sampled mean orientation of the gratings from the interval [1º, 10º], the variability of orientations from the interval [1º,20º], and stimulus contrast from the interval [0.01,1]. In Experiment 2, we sampled the orientation of the gratings from the interval [1º, 45º], the contrast of the dominant grating from the interval [.01,1] and the difference in contrast between the dominant and non-dominant gratings from the interval [.01, *x*] where *x* refers to the contrast of the dominant grating which sets an upper bound on the contrast of the non-dominant grating. In Experiment 3, we sampled the contrast of the Gabor patch from the interval [.01, 1], and noise (in units of standard deviation) from the interval [.01, 2]. Training was validated on a set of 1000 images generated using the same stimulus parameter distributions as the training set.

### Network architectures

We tested three CNN architectures – a 4-layer CNN, VGG-19, and ResNet-50 – on the experiments described above. The networks receive inputs in the form of an image consisting of n x n pixels (n = 90 for Experiment 1 and n = 100 for Experiments 2 and 3) and outputs a binary category label corresponding to the identity of the stimulus (CCW or CW).

The 4-layer CNN model consisted of two convolutional layers (with kernels of size 3 x 3 pixels) paired with two max pooling layers (pooling performed over 2 x 2 pixel windows), one flat layer, and two fully connected layers (consisting of 64 units and 1 unit respectively). A rectified linear unit (ReLu) activation function transformed the outputs of each convolutional layer and the 64-unit fully connected layer, whereas a sigmoid activation function was applied to the output of the final layer.

We also trained two deep CNNs using the standard VGG-19 and ResNet-50 model variants. The VGG-19 model consists of 16 convolutional layers, 3 fully connected layers, 5 max pool layers, and 1 softmax layer. The ResNet-50 model consists of 48 convolutional layers, 1 max pool layer, and 1 average pool layer. The top layer of these networks was modified for binary classification by adding a fully connected layer consisting of a single unit with a sigmoid activation function.

### Training the networks

We trained networks on 10,000 images from each of the three experiments to achieve a classification accuracy > 89% on all tasks. Model performances were assessed on a validation set consisting of 1000 images. The 4-layer CNNs were trained for 25 epochs with a batch size of 32, using the binary cross-entropy loss function and Adam optimizer with a learning rate = 0.001, weight decay = 0 and ε = 10^−8^. As the tasks were relatively simple, to prevent overfitting, we used early stopping with a patience of 10 epochs.

The deep CNNs (VGG-19 and ResNet-50) were trained on these tasks using transfer learning and fine-tuning. We first instantiated the base model pretrained on the ImageNet dataset (provided in Keras Applications at https://keras.io/api/applications/) and froze the model’s weights. The classification layer at the top was excluded to enable feature extraction. Next, we added a global average pooling layer to convert the features extracted from each image into a single vector. Finally, we added a classification head with a single unit to convert these features into binary predictions. To prevent overfitting, we also included a drop-out layer with a drop-out rate of 0.2. Using a base learning rate of 0.001 for the Adam optimizer, we trained this model initially on 10 epochs on binary cross-entropy loss. We found that these networks generally showed poor classification performance (∼60%), and therefore trained them further by unfreezing and fine-tuning the top layers of the network. For fine-tuning, training was continued for a further 10 epochs. The models were fine-tuned on binary cross-entropy loss using a lower learning rate (0.0001) for the RMSprop optimizer. Fine-tuning improved the models’ performances considerably with all models now achieving a classification accuracy of at least 89%.

For each type of network and each experiment, we determined the optimal number of layers to fine-tune by incrementing the number of fine-tuning layers in steps and assessing model performance. We chose the model that gave us the highest accuracy while minimizing the number of layers to fine-tune. For VGG-19, the best models consisted of 8 fine-tuned layers for Experiments 1 and 2 and 5 fine-tuned layers for Experiment 3. For ResNet-50, the best models consisted of 40 fine-tuned layers for Experiments 1 and 2 and 10 fine-tuned layers for Experiment 3.

Finally, to allow for individual differences in learning, we trained 25 instances of each of the three models separately for each experiment using a different random seed to initialize the network’s weights before training.

### Determining stimulus parameters for energy manipulations

To induce confidence-accuracy dissociations, we need to jointly manipulate the signal strength and variability/negative evidence (“energy”) of the stimulus such that the network’s accuracies are matched across conditions. Therefore, we need to determine the stimulus parameters that will allow us to obtain matched network performances across the three conditions. To do so, we first performed a coarse search by fixing the stimulus along the “contrast” dimension for each energy condition (contrast of the Gabor patches for Experiments 1 and 3 and contrast of the dominant grating for Experiment 2) and varying it along the “noise” dimension (variability of orientations for Experiment 1, contrast of the non-dominant grating for Experiment 2 and noise in Experiment 3) in relatively large steps. Next, for each energy level, we determined a range of noise values that gave us a target accuracy between 65-75% and performed a fine-grained search within this range for the parameters that resulted in an accuracy of 70%.

The search yielded stimulus parameter estimates that resulted in matched accuracy levels of 70% across the three energy conditions for each of the three types of networks. Specifically, we obtained the following parameters for the 4-layer CNNs (Experiment 1: contrast = [.2, .25, .3], orientation variance = [7.35°, 21.28°, 27.28°]; Experiment 2: dominant contrast = [0.2, 0.4, 0.6], non-dominant contrast = [0.168, 0.375, 0.575]; Experiment 3: contrast = [.05, .1, .15], noise = [.42, .82, 1.21]), VGG-19 (Experiment 1: contrast = [.4, .5, .6] and orientation variance = [21.42°, 25.28°, 27°]; Experiment 2: dominant contrast = [0.2, 0.4, 0.6] and non-dominant contrast = [0.13, 0.358, 0.56]; Experiment 3: contrast = [.05, .1, .15] and noise = [.32, .54, .715]), and ResNet-50 (Experiment 1: contrast = [.4, .5, .6] and orientation variance = [17°,18.5°, 20°]; Experiment 2: dominant contrast = [0.2, 0.4, 0.6] and non-dominant contrast = [0.13, 0.358, 0.555]; Experiment 3: contrast = [.05, .1, .15] and noise = [.29, .46, .607]). Using these parameters, for each of the three energy levels, we generated stimulus sets consisting of 1000 images to test the CNNs for confidence-accuracy dissociations.

### Behavioral analyses

#### Accuracy and confidence of the networks

The final layer of the network consists of a single unit whose activation (*a*) arises from a sigmoid activation function. The network’s responses (*r*) were generated such that, 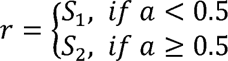 and decision confidence (*c*) was generated as, 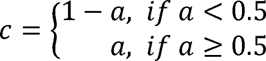 where *a* ϵ [0, 1].

We computed the average accuracy and confidence separately for each of the 25 network instances and for each energy condition.

#### Measures of type-1 and type-2 sensitivity

An observer’s type-1 or perceptual sensitivity (d’) is a measure derived from signal detection theory (SDT) which quantifies the observer’s ability to discriminate between the two stimulus categories (Green, D.M. and Swets, 1966). Type-1 sensitivity (*d’*) is defined as, 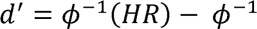(*FAR*) where HR and FAR refer to the observed hit rate and false alarm rates, respectively, when the stimulus category *S*_2_ is treated as the target, and φ^−1^ is the inverse of the cumulative standard normal distribution that transforms cumulative probabilities into z-scores.

Type-2 or metacognitive sensitivity (meta-d’) is a measure derived from SDT-modelling of the observer’s decision and confidence responses which quantifies the information underlying the metacognitive judgement (Maniscalco & Lau, 2012). Intuitively, it can be thought of as a measure of the observer’s ability to distinguish between their own correct and incorrect responses using confidence responses.

### Assessing the networks’ internal activations

To understand the effect of energy manipulations on the internal representations of the network, we studied how the distribution of the network’s activations change with energy levels. Each time an image is presented to the network, it produces an activation in the output layer. We aggregated these activations across all instances of the network separately for all images from each energy condition. We then visualized the distributions of these activations separately for images from each stimulus category using kernel density plots.

To quantify changes in the characteristics of these distributions, we computed two measures – the difference in means of the *S*_1_ and *S*_2_ distributions (which quantifies the separation in evidence between the two categories) and standard deviation of these distributions averaged across the two distributions (which quantifies the degree of uncertainty associated with identity of the stimulus). We computed these measures separately for each network instance and energy condition, and averaged across network instances.

### Simulating the task paradigm for generating meta-d’-d’ dissociations

It has been previously demonstrated that a certain task paradigm can induce dissociations between observers’ meta-d’ and d’ (Maniscalco et al., 2016; Webb et al., 2023). Particularly, in a two-choice task involving discrimination between two stimulus categories (*S*_1_ and *S*_2_), when the contrast of one stimulus category (*S*_1_) is held fixed while the contrast of the other category is allowed to vary across trials (*S*_2_), meta-d’ is found to increase with d’ on trials where the observer responds “*S*_2_ ” and found to decrease with d’ on trials where the observer responds “*S*_1_.”

We simulated this paradigm for Experiments 1-3 by fixing the stimulus contrast for one of the stimulus categories (CCW) and allowing the contrast of the stimuli from the other category (CW) to vary discretely across five levels. Specifically, in Experiment 1, the CCW stimulus was fixed at .225 and the CW-tilted stimuli was varied along the range [.05, .135, .225, .3125, .4]. In Experiment 2, the CCW was fixed at .21 and the CW tilted stimuli was varied along the range [.2, .205, .21, .215, .22]. Finally, in Experiment 3, the CCW was fixed at .1 and the CW tilted stimuli was varied along the range [.05, .075, .1, .125, .15]. The contrast levels of the stimuli were chosen via simulations such that the network’s perceptual sensitivity (d’) spanned a range of meaningful values (1 to 3.5) while avoiding floor or ceiling effects. We generated test sets of 1000 images for each contrast level assumed by *S*_2_ and obtained the decision and confidence responses from our previously trained networks.

We computed d’ and meta-d’ separately for each contrast level of *S*_2_ . For response-specific assessment of meta-d’, we computed meta-d’ separately for trials conditioned on each type of network response (*S*_1_ vs *S*_2_ responses) and contrast level.

### Color discrimination experiment

The task paradigm for the color discrimination experiment was adapted from Desender et al. (2018). This task was previously shown to produce confidence-accuracy dissociations in humans where confidence decreased with stimulus energy levels (Boldt et al., 2017, 2019; Desender et al., 2018; Spence et al., 2016, 2018), in contrast with previous findings where confidence increased with energy levels.

The stimuli (90 x 90 pixels) consisted of an array of eight colored circular patches. Each individual color patch in the array spanned 30 x 30 pixels. The task was to decide whether the average color across the 8 patches was more blue or red (**Figure 6A**). For each image, the mean color across the eight patches was selected from a uniform distribution with a mean color intensity of *c* and an interval of width *v*. The stimulus parameter *c* ϵ [0,1] controlled the intensity of “redness” or “blueness” along a continuous range such that *c* = 1 yielded a completely red patch and *c* = 1 yielded a completely blue patch and values in-between resulted in patches containing a mixture of red and blue with varying proportions of each color. In general, a patch with *c* < .5, contained more red and a patch with *c* > .5 contained more blue. The parameter *v* controlled the variance of color in these patches with higher values of *v* resulting in more variable colors within an array. The energy of the stimulus was manipulated across three levels by simultaneously varying these two features of the array – the color intensity and the spread of color intensity. While increasing the color intensity made it easy to identify the “blueness” or “redness” of the color and made the task easier, increasing the color variance increased the uncertainty regarding the mean color across the patches and made the task harder.

As in our main analyses, we trained 25 instances of the three networks (4-layer CNN, VGG-19 and ResNet-50) using the same procedure outlined above. We determined the stimulus parameters that would allow us to obtain matched network performances across the three energy conditions. The search yielded stimulus parameter estimates that resulted in matched accuracy levels of 70% across the three energy conditions for each of the three types of networks – 4-layer CNNs (color intensity = [.493, .492, .49], color variance = [.4, .494, .626]), VGG-19 and ResNet-50 (contrast = [.4, .3, .2], color variance = [.85, .95, .97]). Using these parameters, for each of the three energy levels, we generated stimulus sets consisting of 1000 images and tested the accuracy and confidence of each of the 25 instances on these images. Finally, as before, we examined the separability and spread of the internal representations of evidence.

### Data and code availability

Codes for training the models and performing all analyses are publicly available at https://osf.io/e5d96/.

## Supporting information

Supplementary analysis

## Acknowledgements

This work was supported by the National Institute of Health (award: R01MH119189) and the Office of Naval Research (award: N00014-20-1-2622).

## References

Aguilar-Lleyda, D., Konishi, M., Sackur, J., & de Gardelle, V. (2021). Confidence can be automatically integrated across two visual decisions. Journal of Experimental Psychology. Human Perception and Performance, 47(2). 10.1037/XHP0000884

Allen, M., Frank, D., Schwarzkopf, D. S., Fardo, F., Winston, J. S., Hauser, T. U., & Rees, G. (2016). Unexpected arousal modulates the influence of sensory noise on confidence. ELife, 5, e18103. 10.7554/eLife.18103

Bang, J. W., Shekhar, M., & Rahnev, D. (2019). Sensory noise increases metacognitive efficiency. Journal of Experimental Psychology: General, 148(3), 437–452. 10.1037/xge0000511

Blauch, N. M., Behrmann, M., & Plaut, D. C. (2021). Computational insights into human perceptual expertise for familiar and unfamiliar face recognition. Cognition, 208, 104341. 10.1016/J.COGNITION.2020.104341

Boldt, A., de Gardelle, V., & Yeung, N. (2017). The impact of evidence reliability on sensitivity and bias in decision confidence. Journal of Experimental Psychology: Human Perception and Performance, 43(8), 1520–1531. 10.1037/xhp0000404

Boldt, A., Schiffer, A. M., Waszak, F., & Yeung, N. (2019). Confidence Predictions Affect Performance Confidence and Neural Preparation in Perceptual Decision Making. Scientific Reports 2019 9:1, 9(1), 1–17. 10.1038/s41598-019-40681-9

Boldt, A., Sun, Y., & Desender, K. (2024). Dis-confirmatory evidence drives confidence. 10.31234/OSF.IO/TSR9Z

Boundy-Singer, Z. M., Ziemba, C. M., & Goris, R. L. T. (2023). Confidence reflects a noisy decision reliability estimate. Nature Human Behaviour, 7(1), 142–154. 10.1038/s41562-022-01464-x

Cao, R., & Yamins, D. (2021). Explanatory models in neuroscience: Part 2 -- constraint-based intelligibility. https://arxiv.org/abs/2104.01489v2

de Gardelle, V., & Mamassian, P. (2015). Weighting Mean and Variability during Confidence Judgments. PLOS ONE, 10(3), e0120870. 10.1371/journal.pone.0120870

De Gardelle, V., & Summerfield, C. (2011). Robust averaging during perceptual judgment. Proceedings of the National Academy of Sciences of the United States of America, 108(32), 13341–13346. 10.1073/PNAS.1104517108/SUPPL_FILE/PNAS.201104517SI.PDF

Desender, K., Boldt, A., & Yeung, N. (2018). Subjective Confidence Predicts Information Seeking in Decision Making. Psychological Science, 29(5), 761–778. 10.1177/0956797617744771

Dobs, K., Martinez, J., Kell, A. J. E., & Kanwisher, N. (2022). Brain-like functional specialization emerges spontaneously in deep neural networks. Science Advances, 8(11). 10.1126/SCIADV.ABL8913

Doerig, A., Sommers, R. P., Seeliger, K., Richards, B., Ismael, J., Lindsay, G. W., Kording, K. P., Konkle, T., van Gerven, M. A. J., Kriegeskorte, N., & Kietzmann, T. C. (2023). The neuroconnectionist research programme. Nature Reviews Neuroscience 2023 24:7, 24(7), 431–450. 10.1038/s41583-023-00705-w

Fetsch, C. R., Kiani, R., Newsome, W. T., & Shadlen, M. N. (2014). Effects of Cortical Microstimulation on Confidence in a Perceptual Decision. Neuron, 83(4), 797–804. 10.1016/j.neuron.2014.07.011

Fleming, S. M., & Daw, N. D. (2017). Self-evaluation of decision-making: A general Bayesian framework for metacognitive computation. Psychological Review, 124(1), 91–114. 10.1037/rev0000045

Fleming, S. M., Maniscalco, B., Ko, Y., Amendi, N., Ro, T., & Lau, H. (2015). Action-Specific Disruption of Perceptual Confidence. Psychological Science, 26(1), 89–98. 10.1177/0956797614557697

Gajdos, T., Fleming, S. M., Saez Garcia, M., Weindel, G., Davranche, K., Garcia, M. S., Weindel, G., Davranche, K., Saez Garcia, M., Weindel, G., & Davranche, K. (2019). Revealing subthreshold motor contributions to perceptual confidence. Neuroscience of Consciousness, 2019(1), niz001. 10.1093/nc/niz001

Gao, Y., Xue, K., Odegaard, B., & Rahnev, D. (2023). Common computations in automatic cue combination and metacognitive confidence reports. BioRxiv. 10.1101/2023.06.07.544029

Gomez-Villa, A., Martin, A., Vazquez-Corral, J., & Bertalmio, M. (2019). Convolutional neural networks can be deceived by visual illusions. Proceedings of the IEEE Computer Society Conference on Computer Vision and Pattern Recognition, 2019-June, 12301–12309. 10.1109/CVPR.2019.01259

Green, D.M. and Swets, J. A. (1966). (1966). Signal Detection Theory and Psychophysics. Wiley, New York. - References - Scientific Research Publish. John Wiley. http://www.scirp.org/(S(lz5mqp453edsnp55rrgjct55))/reference/ReferencesPapers.aspx?ReferenceID=1718728

Green, M. L., Hu, M., Denison, R. N., & Rahnev, D. (2024). Using artificial neural networks to relate external sensory features to internal decisional evidence. 10.31234/OSF.IO/B8AH2

Guggenmos, M. (2022). Reverse engineering of metacognition. ELife, 11. 10.7554/ELIFE.75420

Guo, C., Pleiss, G., Sun, Y., & Weinberger, K. Q. (2017). On Calibration of Modern Neural Networks. 34th International Conference on Machine Learning, ICML 2017, 3, 2130–2143. https://arxiv.org/abs/1706.04599v2

Herce Castañón, S., Moran, R., Ding, J., Egner, T., Bang, D., & Summerfield, C. (2019). Human noise blindness drives suboptimal cognitive inference. Nature Communications, 10(1). 10.1038/S41467-019-09330-7

Kell, A. J., & McDermott, J. H. (2019). Deep neural network models of sensory systems: windows onto the role of task constraints. Current Opinion in Neurobiology, 55, 121–132. 10.1016/J.CONB.2019.02.003

Kell, A. J., Yamins, D. L. K., Shook, E. N., Norman-Haignere, S. V., & McDermott, J. H. (2018). A Task-Optimized Neural Network Replicates Human Auditory Behavior, Predicts Brain Responses, and Reveals a Cortical Processing Hierarchy. Neuron, 98(3), 630–644.e16. 10.1016/J.NEURON.2018.03.044

Koizumi, A., Maniscalco, B., & Lau, H. (2015). Does perceptual confidence facilitate cognitive control? Attention, Perception & Psychophysics, 77(4), 1295–1306. 10.3758/S13414-015-0843-3

Koriat, A. (2006). Metacognition and consciousness. The Cambridge Handbook of Consciousness, 3(2), 289–326. 10.1017/CBO9780511816789.012

Li, H. H., & Ma, W. J. (2020). Confidence reports in decision-making with multiple alternatives violate the Bayesian confidence hypothesis. Nature Communications, 11(1), 1–11. 10.1038/s41467-020-15581-6

Mamassian, P. (2016). Visual Confidence. 10.1146/Annurev-Vision-111815-114630, 2, 459–481. 10.1146/ANNUREV-VISION-111815-114630

Mamassian, P., & de Gardelle, V. (2021). Modeling perceptual confidence and the confidence forced-choice paradigm. Psychological Review. 10.1037/REV0000312

Maniscalco, B., & Lau, H. (2012). A signal detection theoretic approach for estimating metacognitive sensitivity from confidence ratings. Consciousness and Cognition, 21(1), 422–430. 10.1016/j.concog.2011.09.021

Maniscalco, B., & Lau, H. (2016). The signal processing architecture underlying subjective reports of sensory awareness. Neuroscience of Consciousness, 2016(1), niw002. 10.1093/nc/niw002

Maniscalco, B., Peters, M. A. K., & Lau, H. (2016). Heuristic use of perceptual evidence leads to dissociation between performance and metacognitive sensitivity. *Attention, Perception*, & Psychophysics, 78(3), 923–937. 10.3758/s13414-016-1059-x

Metcalfe, J., & Shimamura, A. P. (1994). Metcalfe, J., & Shimamura, A. P. (1994). Metacognition: Knowing about Knowing. Cambridge, MA: MIT Press. - References - Scientific Research Publish. In MIT Press. http://www.scirp.org/(S(351jmbntvnsjt1aadkposzje))/reference/ReferencesPapers.aspx?ReferenceID=1738390

Minderer, M., Djolonga, J., Romijnders, R., Hubis, F., Zhai, X., Houlsby, N., Tran, D., & Lucic, M. (2021). Revisiting the Calibration of Modern Neural Networks. Advances in Neural Information Processing Systems, 19, 15682–15694. https://arxiv.org/abs/2106.07998v2

Morales, J., Solovey, G., Maniscalco, B., Rahnev, D., de Lange, F. P., & Lau, H. (2015). Low attention impairs optimal incorporation of prior knowledge in perceptual decisions. *Attention*, Perception, and Psychophysics, 77(6), 2021–2036. 10.3758/S13414-015-0897-2/FIGURES/12

Odegaard, B., Grimaldi, P., Cho, S. H., Peters, M. A. K., Lau, H., & Basso, M. A. (2018). Superior colliculus neuronal ensemble activity signals optimal rather than subjective confidence. Proceedings of the National Academy of Sciences of the United States of America, 115(7), E1588–E1597. 10.1073/PNAS.1711628115/SUPPL_FILE/PNAS.201711628SI.PDF

Peters, M. A. K., Thesen, T., Ko, Y. D., Maniscalco, B., Carlson, C., Davidson, M., Doyle, W., Kuzniecky, R., Devinsky, O., Halgren, E., & Lau, H. (2017). Perceptual confidence neglects decision-incongruent evidence in the brain. Nature Human Behaviour, 1(7), 0139. 10.1038/s41562-017-0139

Rahnev, D., Koizumi, A., McCurdy, L. Y., Esposito, M. D., Lau, H., D’Esposito, M., & Lau, H. (2015). Confidence Leak in Perceptual Decision Making. Psychological Science, 26(11), 1664–1680. 10.1177/0956797615595037

Rahnev, D., Kok, P., Munneke, M., Bahdo, L., De Lange, F. P., & Lau, H. (2013). Continuous theta burst transcranial magnetic stimulation reduces resting state connectivity between visual areas. Journal of Neurophysiology, 110(8), 1811–1821. 10.1152/jn.00209.2013

Rahnev, D., Maniscalco, B., Graves, T., Huang, E., De Lange, F. P., & Lau, H. (2011). Attention induces conservative subjective biases in visual perception. Nature Neuroscience, 14(12), 1513–1515. 10.1038/nn.2948

Rahnev, D., Maniscalco, B., Luber, B., Lau, H., & Lisanby, S. H. (2012). Direct injection of noise to the visual cortex decreases accuracy but increases decision confidence. Journal of Neurophysiology, 107(6), 1556–1563. 10.1152/jn.00985.2011

Rahnev, D., Nee, D. E., Riddle, J., Larson, A. S., D’Esposito, M., & D’Esposito, M. (2016). Causal evidence for frontal cortex organization for perceptual decision making. Proceedings of the National Academy of Sciences, 113(20), 201522551. 10.1073/pnas.1522551113

Rausch, M., Hellmann, S., & Zehetleitner, M. (2018). Confidence in masked orientation judgments is informed by both evidence and visibility. *Attention*, Perception, and Psychophysics, 80(1), 134–154. 10.3758/s13414-017-1431-5

Rausch, M., Zehetleitner, M., Steinhauser, M., & Maier, M. E. (2020). Cognitive modelling reveals distinct electrophysiological markers of decision confidence and error monitoring. NeuroImage, 218, 116963. 10.1016/j.neuroimage.2020.116963

Richards, B. A., Lillicrap, T. P., Beaudoin, P., Bengio, Y., Bogacz, R., Christensen, A., Clopath, C., Costa, R. P., de Berker, A., Ganguli, S., Gillon, C. J., Hafner, D., Kepecs, A., Kriegeskorte, N., Latham, P., Lindsay, G. W., Miller, K. D., Naud, R., Pack, C. C., … Kording, K. P. (2019). A deep learning framework for neuroscience. Nature Neuroscience 2019 22:11, 22(11), 1761–1770. 10.1038/s41593-019-0520-2

Rounis, E., Maniscalco, B., Rothwell, J. C., Passingham, R. E., & Lau, H. (2010). Theta-burst transcranial magnetic stimulation to the prefrontal cortex impairs metacognitive visual awareness. Cognitive Neuroscience, 1(3), 165–175. 10.1080/17588921003632529

Samaha, J., Barrett, J. J., Sheldon, A. D., LaRocque, J. J., & Postle, B. R. (2016). Dissociating perceptual confidence from discrimination accuracy reveals no influence of metacognitive awareness on working memory. Frontiers in Psychology, 7(JUN), 851. 10.3389/FPSYG.2016.00851/BIBTEX

Shekhar, M., & Rahnev, D. (2018). Distinguishing the Roles of Dorsolateral and Anterior PFC in Visual Metacognition. The Journal of Neuroscience, 38(22), 5078–5087. 10.1523/JNEUROSCI.3484-17.2018

Shekhar, M., & Rahnev, D. (2021). The nature of metacognitive inefficiency in perceptual decision making. Psychological Review, 128(1), 45–70. 10.1037/rev0000249

Shekhar, M., & Rahnev, D. (2023). How do humans give confidence? A comprehensive comparison of process models of perceptual metacognition. 10.31234/OSF.IO/CWRNT

Spence, M. L., Dux, P. E., & Arnold, D. H. (2016). Computations underlying confidence in visual perception. Journal of Experimental Psychology: Human Perception and Performance, 42(5), 671–682. 10.1037/xhp0000179

Spence, M. L., Mattingley, J. B., & Dux, P. E. (2018). Uncertainty information that is irrelevant for report impacts confidence judgments. Journal of Experimental Psychology: Human Perception and Performance, 44(12), 1981–1994. 10.1037/xhp0000584

Webb, T. W., Miyoshi, K., So, T. Y., Rajananda, S., & Lau, H. (2023). Natural statistics support a rational account of confidence biases. Nature Communications 2023 14:1, 14(1), 1–18. 10.1038/s41467-023-39737-2

Wichmann, F. A., & Geirhos, R. (2023). Are Deep Neural Networks Adequate Behavioral Models of Human Visual Perception? Annual Review of Vision Science, 9, 501–524. 10.1146/ANNUREV-VISION-120522-031739

Wilimzig, C., Tsuchiya, N., Fahle, M., Einhäuser, W., & Koch, C. (2008). Spatial attention increases performance but not subjective confidence in a discrimination task. Journal of Vision, 8(5), 7. 10.1167/8.5.7

Wokke, M. E., Achoui, D., & Cleeremans, A. (2020). Action information contributes to metacognitive decision-making. Scientific Reports, 10(1), 1–15. 10.1038/s41598-020-60382-y

Xue, K., Zheng, Y., Rafiei, F., & Rahnev, D. (2023). The timing of confidence computations in human prefrontal cortex. BioRxiv : The Preprint Server for Biology. 10.1101/2023.03.21.533662

Zylberberg, A., Barttfeld, P., & Sigman, M. (2012). The construction of confidence in a perceptual decision. Frontiers in Integrative Neuroscience, 6, 79. 10.3389/fnint.2012.00079

Zylberberg, A., Fetsch, C. R., & Shadlen, M. N. (2016). The influence of evidence volatility on choice, reaction time and confidence in a perceptual decision. ELife, 5(OCTOBER2016), e17688. 10.7554/eLife.17688

Zylberberg, A., Roelfsema, P. R., & Sigman, M. (2014). Variance misperception explains illusions of confidence in simple perceptual decisions. Consciousness and Cognition, 27, 246–253. 10.1016/j.concog.2014.05.012

