## Supplementary analysis for "Human-like dissociations between confidence and accuracy in convolutional neural networks"

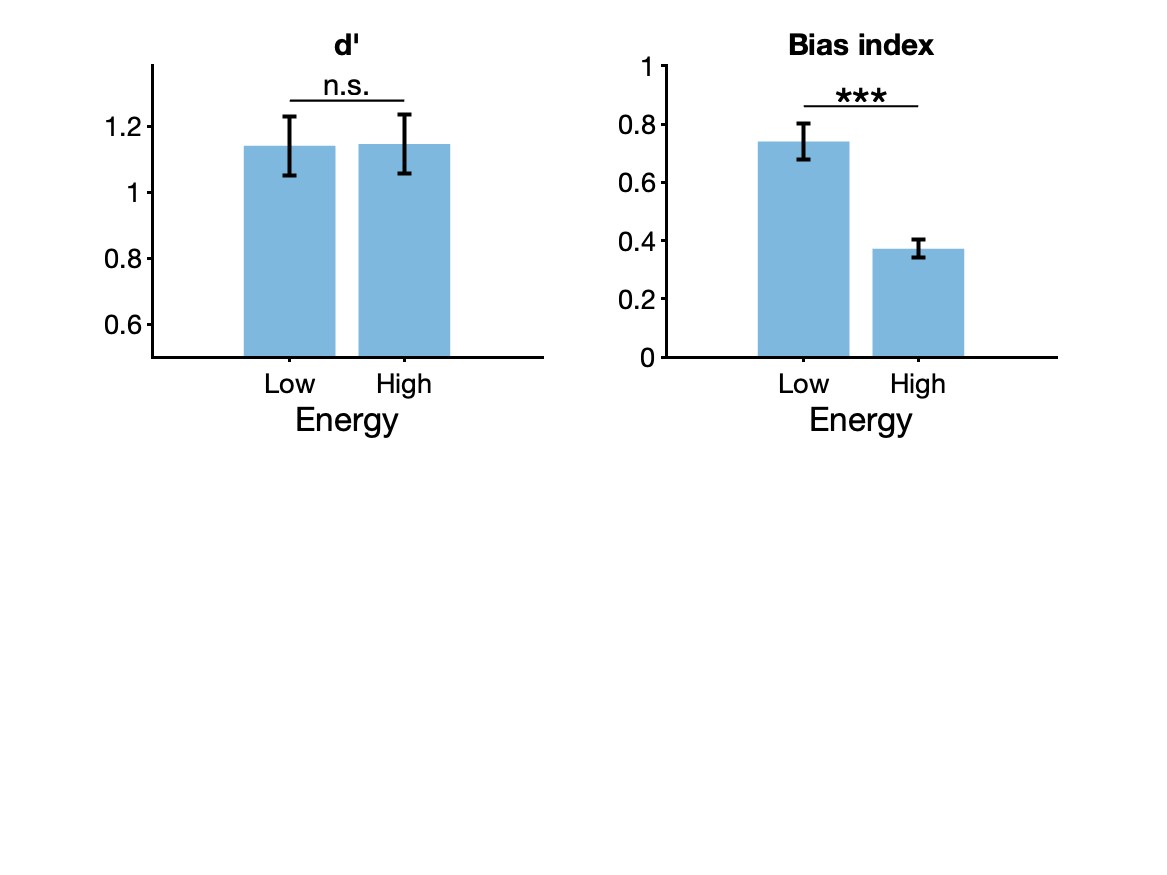
**Supplementary Figure 1. The signal-and-variance-increase hypothesis can explain the effect of base-rate changes on choice behavior during energy manipulations.** Herce Castañón et al. (2019) showed that when subjects are asked to judge the mean orientation of an array of Gabor patches, they exhibit a range of suboptimal behaviors. Specifically, Herce Castañón et al. tested how energy manipulation interact with changes in stimulus base rates by measuring the shifts in subjects’ decision criteria in response to changes in stimulus frequencies for each energy condition. The location of the decision criteria was computed from the SDT-based measure of response bias ($c$). The shift in criteria in response to changes in stimulus base rates was quantified as $c_{S_{1}}-c_{S_{2}}$where $c_{S_{i}}$refers to the criterion in the condition where stimulus $i$is more frequent. Herce Castañón et al. found that subjects showed significantly smaller shifts in their criterion in response to base rate changes in the high-energy condition, compared to the low-energy condition. They explained these effects by proposing that observers are blind to the noise arising from their own cognitive computations, thus resulting in failure to account for the higher levels of uncertainty arising from high-energy stimuli. Here, we show via simulations that the signal-and-variance-increase hypothesis can also explain their observed effects. We simulated the effect of stimulus energy manipulations in an SDT model where higher stimulus energy led to increase in signal ($\mu$) as well as variance ($\sigma$) of the stimulus evidence distributions. We generated 50 simulations by sampling individual SDT parameters: the signal, $\mu$, and the decision criterion, $c,$ from Gaussian distributions. Specifically, in the low-energy condition $\mu_{low}\sim N\left( 1, .5 \right)$ and in the high-energy condition $\mu_{high}=N\left( 2, 1 \right)$, such that the two distributions would lead to equal sensitivity but the high-energy condition features higher variance of the internal distributions. Similarly, the decision criterion for individual simulations was sampled from $c\sim N\left( 0, .25 \right)$. For the base-rate manipulations, we assumed that an increase in probability of observing the stimulus class $S_{1}$ would shift the criterion by an amount $+c_{shift}$to allow more frequent $S_{1}$ responses. Similarly, increase in probability of $S_{2}$ stimuli would lead to a criterion shift of $-c_{shift}$. We allowed individual variability in criterion shifts by sampling $c_{shift} \sim N\left( .4,.2 \right).$ Critically, we assumed that these criteria were fixed across the two energy conditions. Using these parameters, we generated data for 10,000 trials from each of the 50 simulations and computed the SDT-derived measures of stimulus sensitivity ($d_{obs}^{'}$) and response bias ($c_{obs})$ from these data separately for each energy condition and base-rate condition. As done by Herce Castañón et al. (2019), we computed the bias index as the difference in $c_{obs}$ between the two base-rate conditions. The plots show the average $d_{obs}^{'}$ and bias index across all simulations, separately for the low- and high-energy conditions. Our findings replicate the original observations that in spite of matched performance between the two energy conditions ($\Delta d^{'}=.0024, t\left( 49 \right)=-.684, p=.50)$, observers appear to shift their decision criterion less in favor of the more probable stimulus in the high-energy condition ($\Delta bias index=.39, t\left( 49 \right)=13.66, p<.0001)$. These findings show that the signal-and-variance-increase hypothesis can indeed capture the suboptimal behaviors attributed to the noise-blindness mechanism. ***p<0.001; n.s., not significant.
